## Supplementary Information for "Structure and mechanism of biosynthesis of *Streptococcus mutans* cell wall polysaccharide"

**Supplementary Table 1.** Activity of membrane fractions of *E. coli* strains, carrying empty expression vector, vector with *sccN* or vector with *sccP* in presence of Und-P and UDP-[^3^H]Glc ^a^

| **Gene expressed** | **Activity,**  **pmol/mg** |
| --- | --- |
| None | 0.4 |
| *sccN* | 603.6 |
| *sccP* | 278.0 |

^a^ Reaction mixtures contained 50 mM Tris-Cl, pH 7.4, 20 mM MgCl_2_, 2 mM ATP, 2 mM sodium orthovanadate, 10 mM 2-mercaptoethanol, 15 μM Und-P, dispersed ultrasonically in 1 % CHAPS, 0.35 % CHAPS (final concentration in the reaction), 5 μM UDP-Glc (1600 cpm/pmol) and 20-60 μg solubilized *E  coli* membrane proteins (expressing *sccN* or *sccP* on a plasmid or carrying empty expression vector) in a total volume of 0.02 mL. Following 3 min at 37 °C reactions were stopped by the addition of 2 mL CHCl_3_/CH_3_OH and the [^3^H]glucolipid product isolated as described in Methods.

**Supplementary Table 2**. Glycosyl linkages detected in rhamnopolysaccharides from various strains of *S. mutans* determined as partially methylated alditol acetates ^a^

| **Glycosyl**  **linkage** | **WT** | | | **Mean±SD** | **Δ*sccN*** | | | **Mean±SD** | **Δ*sccP*** | | | **Mean±SD** | | **Δ*sccQ*** | | | **Mean±SD** | **Δ*sccM*** | | | **Mean±SD** |
| --- | --- | --- | --- | --- | --- | --- | --- | --- | --- | --- | --- | --- | --- | --- | --- | --- | --- | --- | --- | --- | --- |
|  | **mol/mol** | | |  | **mol/mol** | | |  | **mol/mol** | | |  |  | **mol/mol** | | |  | **mol/mol** | | |  |
| t-Rha*p* | 0.1 | 0.3 | 0.2 | 0.2±0.1 | 0.3 | 0.8 | 0.4 | 0.5±0.3 | 0.1 | 0.4 | 0.2 | | 0.2±0.2 | 0.3 | 0.7 | 0.4 | 0.4±0.2 | 0.2 | 0.5 | 0.3 | 0.3±0.2 |
| 2-Rha*p* | 20.9 | 19.2 | 22.9 | 21±1.9 | 22.8 | 21.7 | 23.5 | 22.7±0.9 | 19.7 | 17.3 | 22.5 | | 19.9±2.6 | 22.0 | 20.2 | 23.3 | 21.8±1.6 | 16.9 | 16.8 | 17.9 | 17.2±0.6 |
| 3-Rha*p* | 10.4 | 10.2 | 9.2 | 9.9±0.6 | 23.5 | 21.9 | 22.2 | 22.5±0.8 | 9.8 | 10.5 | 8.9 | | 9.7±0.8 | 19.2 | 18.0 | 18.3 | 18.5±0.6 | 24.3 | 21.5 | 22.9 | 22.9±1.4 |
| 3,4-Rha*p* | 0.2 | 0.3 | 0.1 | 0.2±0.1 | 3.2 | 4.8 | 3.4 | 3.8±0.9 | 0.0 | 0.1 | 0.0 | | 0.1±0.1 | 0.1 | 0.2 | 0.0 | 0.1±0.1 | 1.8 | 2.5 | 1.8 | 2.1±0.4 |
| 2,3-Rha*p* | 14.7 | 14.7 | 14.3 | 14.6±0.3 | 0.2 | 0.5 | 0.3 | 0.3±0.2 | 16.8 | 16.6 | 15.4 | | 16.3±0.8 | 7.8 | 9.4 | 7.1 | 8.1±1.2 | 0.2 | 0.4 | 0.3 | 0.3±0.1 |
| 2,4-Rha*p* | 1.8 | 2.5 | 1.6 | 2±0.5 | 0.1 | 0.3 | 0.2 | 0.2±0.1 | 3.4 | 4.8 | 2.9 | | 3.7±1.0 | 0.7 | 1.5 | 0.8 | 1±0.4 | 6.7 | 8.2 | 6.8 | 7.2±0.9 |
| 2,3,4-Rha*p* | 1.9 | 2.7 | 1.8 | 2.1±0.5 | n.d.* | n.d.* | n.d.* |  | n.d.* | 0.1 | n.d.* | |  | n.d.* | 0.1 | n.d.* |  | n.d.* | 0.1 | n.d.* |  |
| t-Glc*p* | 20.5 | 17.2 | 19.2 | 19±1.7 | 3.4 | 5.2 | 3.3 | 4±1.1 | 20.0 | 17.5 | 19.1 | | 18.8±1.3 | 8.3 | 9.2 | 6.9 | 8.1±1.1 | 8.6 | 9.7 | 7.7 | 8.7±1.0 |
| t-Glc*p*NAc | 0.1 | 0.6 | 0.3 | 0.3±0.2 | 0.1 | 0.5 | 0.4 | 0.3±0.2 | 0.1 | 1.0 | 0.3 | | 0.5±0.5 | 0.1 | 0.7 | 0.3 | 0.4±0.3 | 0.1 | 0.4 | 0.3 | 0.3±0.2 |
| 4-Glc*p*NAcOl | 0.3 | 0.8 | 0.4 | 0.5±0.3 | 0.2 | 0.8 | 0.5 | 0.5±0.3 | 0.2 | 1.2 | 0.3 | | 0.6±0.6 | 0.3 | 0.9 | 0.3 | 0.5±0.4 | 0.1 | 0.6 | 0.4 | 0.4±0.2 |

^a^ Partially methylated alditol acetates were prepared from purified rhamnopolysacchrides, detected by GC-MS and normalized to a polymer containing an average of 50 Rha units, as described in Methods. *n.d. indicates none detected. Moles of each linkage type per molecule were calculated by multiplying the mol % for each component sugar by the relative area percent for each linkage type found in the gas chromatogram and then normalized to a hypothetical polymer containing 50 Rha units. Individual values of each linkage type from three separate analyses, as well as average and standard deviation calculations, are shown.

**Supplementary Table 3**. Cell size analysis ^a^

| ***S. mutans strain*** | **Cell #** | **Cell length, μm±SD** | **Cell width, μm±SD** |
| --- | --- | --- | --- |
| WT | 144 | 0.86±0.11 | 0.63±0.10 |
| Δ*sccQ* | 198 | 0.76±0.10 | 0.66±0.07 |

^a^ DIC images were used to determine cell sizes of *S. mutans* strains. Cell size (μm) was measured by ImageJ, ObjectJ plugin, and the results were analyzed by GraphPad Prism 9.3. using unpaired two-tailed *t*-test with Welch correction. Total number of cells was n = 144 for WT and n = 198 for Δ*sccQ*. Values are reported with standard deviation.

**Supplementary Table 4.** Bacterial strains and plasmids

| **Strain or plasmid** | **Description ^a^** | **Reference** |
| --- | --- | --- |
| ***Streptococcus mutans*** | | |
| Xc | Serotype *c* strain, wild-type (WT) | ^1^ |
| Δ*sccH* | *sccH* deletion mutant (has a nonpolar erythromycin resistance cassette inserted in *sccH*), Erm^R^ | ^2^ |
| Δ*sccH:*p*sccH* | Δ*sccH* is complemented with p*sccH* carrying WT *sccH,* Erm^R^, Cam^R^ | ^2^ |
| Δ*sccN* | *sccN* deletion mutant (has a nonpolar spectinomycin resistance cassette inserted in *sccN*), Spec^R^ | ^3^ |
| Δ*sccN:*p*sccN* | Δ*sccN* is complemented with p*sccN* carrying WT *sccN,* Spec^R^, Cam^R^ | ^3^ |
| Δ*sccN:*p*gacHIJKL* | Δ*sccN* is complemented with p*gacHIJKL*. Spec^R^, Cam^R^ | ^3^ |
| Δ*sccP* | *sccP* deletion mutant (has a nonpolar spectinomycin resistance cassette inserted in *sccP*), Spec^R^ | ^3^ |
| Δ*sccM* | *sccM* deletion mutant (has a nonpolar spectinomycin resistance cassette inserted in *sccM*), Spec^R^ | This study |
| Δ*sccQ* | *sccQ* deletion mutant (has a nonpolar spectinomycin resistance cassette inserted in *sccQ*), Spec^R^ | This study |
| Δ*sccN*Δ*sccP* | *sccN sccP* double-gene deletion mutant (has nonpolar spectinomycin and erythromycin resistance cassettes inserted in *sccN* and *sccP,* respectively), Spec^R^ Erm^R^ | ^3^ |
| Δ*sccM*Δ*sccQ* | *sccM sccQ* double-gene deletion mutant (has nonpolar erythromycin and spectinomycin resistance cassettes inserted in *sccM* and *sccQ,* respectively), Spec^R^ Erm^R^ | This study |
| Δ*sccN*Δ*sccQ* | *sccN sccQ* double-gene deletion mutant (has nonpolar kanamycin and spectinomycin resistance cassettes inserted in *sccN* and *sccP,* respectively), Kan^R^ Spec^R^ | This study |
| Δ*sccM*Δ*sccN* | *sccM sccN* double-gene deletion mutant (has nonpolar erythromycin and kanamycin resistance cassettes inserted in *sccM* and *sccN,* respectively), Erm^R^ Kan^R^ | This study |
| Δ*sccM*Δ*sccP* | *sccM sccP* double-gene deletion mutant (has nonpolar erythromycin and spectinomycin resistance cassettes inserted in *sccM* and *sccP,* respectively), Erm^R^ Spec^R^ | This study |
| ***Escherichia coli*** | | |
| DH5α | *E. coli* cells used for cloning | Invitrogen |
| JW2347 | *E. coli* K-12 strain BW25113 ^4^ with a deletion of the *gtrB* gene ^5^ | ^6^ |
| ***Plasmids*** |  |  |
| pBAD33_SccN | A pBAD33 derived plasmid expressing *sccN,* Amp^R^, Cam^R^ | This study |
| pBAD33_SccP | A pBAD33 derived plasmid expressing *sccP,* Amp^R^, Cam^R^ | This study |
| pLR16T | Vector encoding spectinomycin resistance cassette. Spec^R^ | ^7^ |
| pOSKAR | Vector encoding kanamycin resistance cassette. Kan^R^ | ^8^ |
| pHY304 | Vector encoding erythromycin resistance cassette. Erm^R^ | ^9^ |

^a^ Antibiotic resistance markers: Erm^R^, erythromycin; Kan^R^, kanamycin; Spec^R^, spectinomycin; Cam^R^, chloramphenicol, Amp^R^, ampicillin

**Supplementary Table 5.** Primers used for construction of bacterial mutants and plasmids

| Primer | Sequence ^a,b^ | Genetic manipulations |
| --- | --- | --- |
| Smu.833-f | GGTTCTGACAGTCGTCTCTC | *sccN* deletion with a nonpolar kanamycin resistance cassette |
| Kan-Smu.833-r1 | **CAGTATTTAAAGATACC**GGTTTCTTCCTCATTATAAC |  |
| Smu.833-Kan-f1 | TAATGAGGAAGAAACC**GGTATCTTTAAATACTGTAG** |  |
| Kan-Smu.833-f2 | **TGAATTGTTTTAGTAC**GATTTACAGGATCCGCCAG |  |
| Smu.833-Kan-r2 | CGGATCCTGTAAATC**GTACTAAAACAATTCATCCAG** |  |
| Smu.833-r | GCAACAAAATTTAGAATCAACAAC |  |
| SccM-f | GTGGGACTTAATGTTAGTG | *sccM* deletion with a nonpolar spectinomycin resistance cassette |
| Spec-SccM -r1 | **CACTATTTTGGTCGAC**CAATAGGCGGTAATGATTC |  |
| SccM-Spec-f1 | GAATCATTACCGCCTATTG**GTCGACCAAAATAGTGAGGAGG** |  |
| Spec-SccM-f2 | **AAAATTATAAGGATCC**GAGATCAAACGATCTTTGCG |  |
| SccM-Spec-r2 | CGCAAAGATCGTTTGATCTC**GGATCCTTATAATTTTTTTAATCTG** |  |
| SccM-r | CTGATACCTAACTTATTTATAATG |  |
| SccM-f | GTGGGACTTAATGTTAGTG | *sccM* deletion with a nonpolar erythromycin resistance cassette |
| Erm-SccM-r1 | **CATCTAATTTAACTTCAATTCC**CAATAGGCGGTAATGATTC |  |
| SccM-Erm-f1 | GAATCATTACCGCCTATTG**GGAATTGAAGTTAAATTAGATG** |  |
| Erm-SccM-f2 | **CGGGAGGAAATAATTCTATG**GAGATCAAACGATCTTTGCG |  |
| SccM-Erm -r2 | CGCAAAGATCGTTTGATCTC**CATAGAATTATTTCCTCCCG** |  |
| SccM-r | CTGATACCTAACTTATTTATAATG |  |
| SccQ-f | GTTAATCACCTTTACCAAGG | *sccQ* deletion with a nonpolar spectinomycin resistance cassette |
| Spec-SccQ-r1 | **CACTATTTTGGTCGAC**GGCTATTCGAAAAATTCCAATG |  |
| SccQ-Spec-f1 | TTTTTCGAATAGCC**GTCGACCAAAATAGTGAGGAGG** |  |
| Spec-SccQ-f2 | **AAAATTATAAGGATCC**GCTATTTCAATTGCTTCAGG |  |
| SccQ-Spec-r2 | GCAATTGAAATAGC**GGATCCTTATAATTTTTTTAATCTG** |  |
| SccQ-r | GCTAATTCGAAAGCTTTTCG |  |
| SccQ-f | GTTAATCACCTTTACCAAGG | *sccQ* deletion with a nonpolar erythromycin resistance cassette |
| Erm-SccQ-r1 | **CATCTAATTTAACTTCAATTCC**GGCTATTCGAAAAATTCCAATG |  |
| SccQ-Erm-f1 | TTTTTCGAATAGCC**GGAATTGAAGTTAAATTAGATG** |  |
| Erm-SccQ-f2 | **CGGGAGGAAATAATTCTATG**GCTATTTCAATTGCTTCAGG |  |
| SccQ-Erm -r2 | GCAATTGAAATAGC**CATAGAATTATTTCCTCCCG** |  |
| SccQ-r | GCTAATTCGAAAGCTTTTCG |  |
| sccM-check-f | GATTCTTTGACATTCGATAAATC | Verification of Δ*sccM* |
| sccM-check-r | GATGTCCTGACATAAACATCG |  |
| sccQcheck-f | GAGGGAGATTGGTCTAATTG | Verification of Δ*sccQ* |
| sccQcheck-r | CAGCTGAGTTAGAGCAAGAAAAAG |  |
| sccN-XbaI-f | GCGACTCTAGACCAATTATTAATTTTCAAGG | Construction of pBAD33_SccN |
| sccN-HindIII-r | CGCGCAAGCTTCCTATAGCCTTTATCCTTTTTC |  |
| SccP-XbaI-f | GCGACTCTAGAAGGAGAATTTATACTATGACAGAG | Construction of  pBAD33_SccP |
| SccP-SalI-r | CGCTGCGTCGACCCTAAAAACTATTTACGGCC |  |

^a^ Restriction sites are underlined.

^b^ Extensions complementary to the antibiotic resistance cassettes are in bold.

**Supplementary Table 6.** Spontaneous mutations detected in the mutants of *S. mutans* Xc by whole-genome sequencing

| **Strain** | **Target insertion of antibiotic resistance cassette^a^** | **Spontaneous mutations^a^** | **Annotation** |
| --- | --- | --- | --- |
| ∆*sccH* | Smu.831 | n.d.^b^ |  |
| ∆*sccM* | Smu.832 | n.d. |  |
| ∆*sccN* | Smu.833 | n.d. |  |
| ∆*sccP* | Smu.834 | Smu.82  V113I (GTT→ATT) | DnaK |
|  |  | Smu.1237c  D12E (TAT→TGT) | Nuclear transport factor 2 family protein |
| ∆*sccQ* | Smu.835 | n.d. |  |
| ∆*sccM*∆*sccN* | Smu.832  Smu.833 | n.d. |  |
| ∆*sccM*∆*sccQ* | Smu.832  Smu.835 | Smu.834  Y37C (TAT→TGT) | SccP |
| ∆*sccN*∆*sccP* | Smu.833  Smu.834 | n.d. |  |
| *∆sccN*∆*sccQ* | Smu.833  Smu.835 | n.d. |  |
| ∆*sccM*∆*sccP* | Smu.832  Smu.834 | Smu.82 V113I (GTT→ATT) | DnaK |
|  |  | Smu.1237c  D12E (TAT→TGT) | Nuclear transport factor 2 family protein |

^a^ corresponds to genes in *S. mutans* UA159 (GenBank: AE014133.2)

^b^ n.d. indicates none detected.

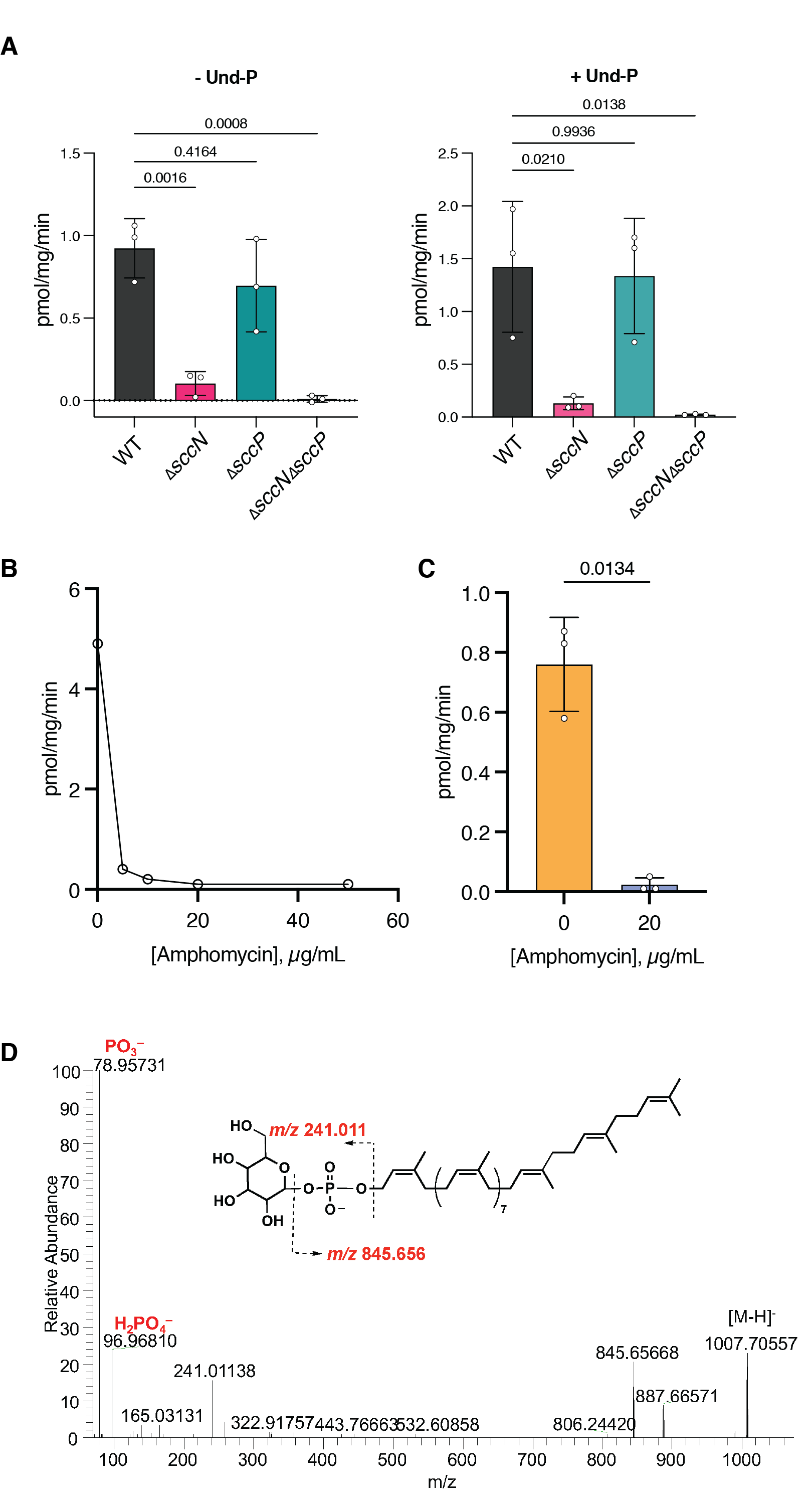

**Supplementary Fig. 1. SccN and SccP synthesize Glc-P-Und**

(**a**) Membrane fractions from various *S.* *mutans* strains were assayed for Glc-P-Und synthase activity in the presence and absence of 20 μM Und-P, added as a sonicated dispersion in 1 % CHAPS, as described in Methods. The results are averages from three independent membrane preparations ± S.D. Significant differences were determined with one-way ANOVA with Tukey’s multiple comparison test. (**b**) Concentration-dependent inhibition of Glc-P-Und synthesis by the addition of amphomycin. Membrane fractions from *S.* *mutans* Xc were incubated with UDP-[^3^H]Glc and the indicated concentrations of amphomycin and assayed for the formation of [^3^H]Glc-P-Und as described in Methods, except that 1 mM CaCl_2_ was included to facilitate the function of the antibiotic. (**c**) Membrane fractions from three independent isolates of *S.* *mutans* Xc were tested, *in vitro*, for Glc-P-Und synthase activity in the presence and absence of 20 μg/mL amphomycin. Two-tailed unpaired t-test with Welch’s correction was used to determine statistical significance. (**d**) ESI-MS analysis of compound(s) co-migrating with synthetic [^3^H]Glc-P-Und purified from the *S. mutans* membrane fraction by preparative TLC.

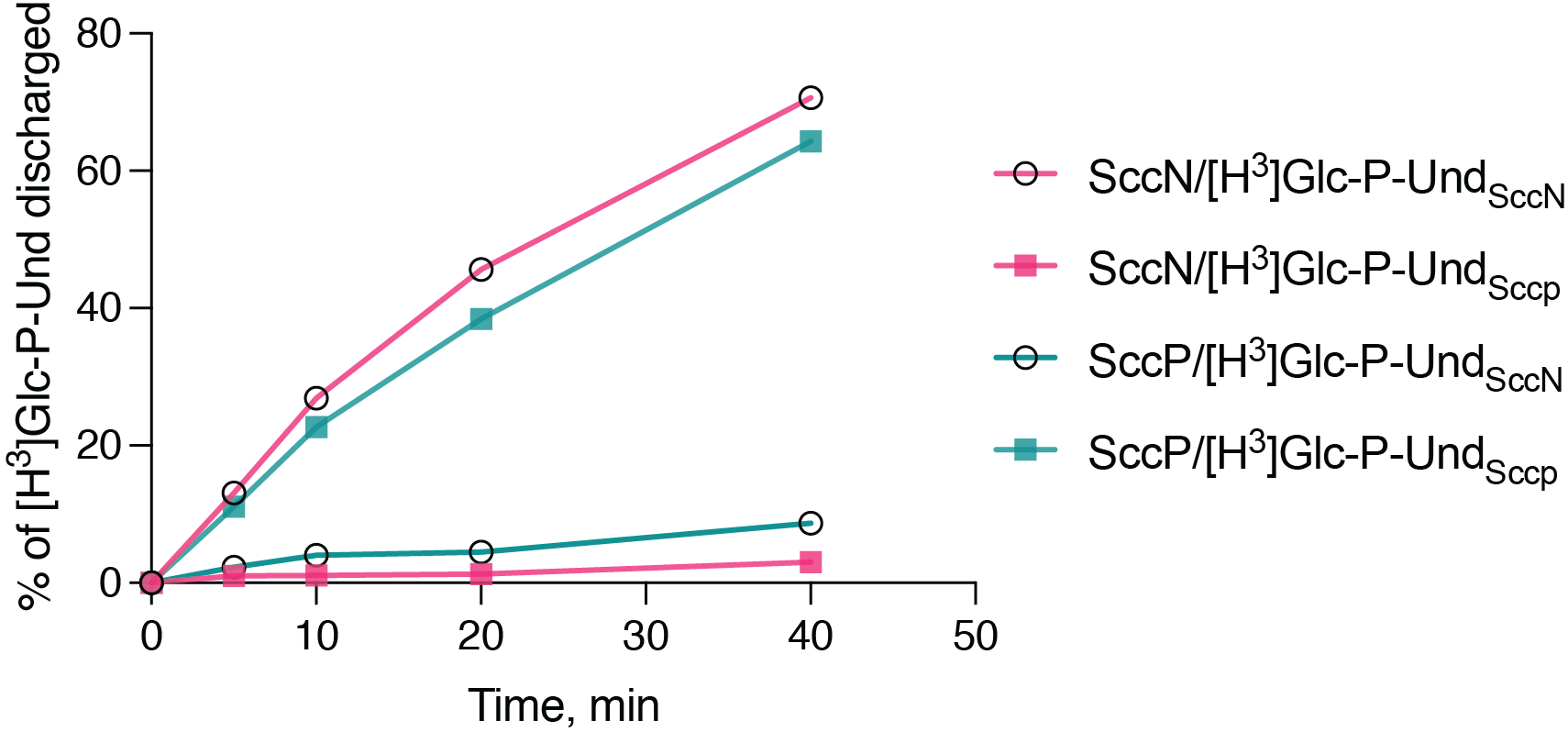

**Supplementary Fig. 2. Time-dependent reverse reactions of SccN and SccP to reform UDP-Glc from their respective enzymatic products and UDP**

[^3^H]Glc-P-Und_SccN_, synthesized by SccN, and [^3^H]Glc-P-Und_SccP_, synthesized by SccP, were tested as substrates in the discharge reactions, containing soluble, partially-purified SccN or SccP, as described in Methods. The experiment was performed independently three times and yielded the same results.

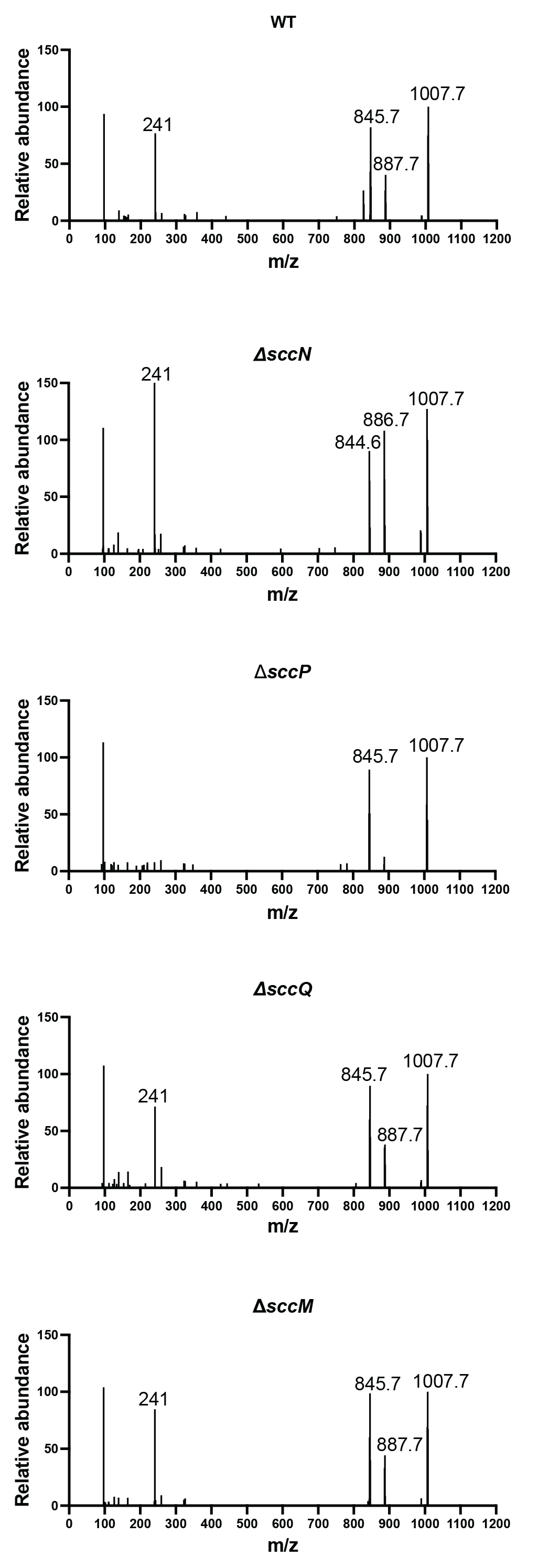

**Supplementary Fig. 3. ESI-MS/MS analysis of Glc-P-Unds purified from *S. mutans* WT, Δ*sccN*, Δ*sccP*, Δ*sccM* and Δ*sccQ* strains by preparative TLC**

Phospholipids were extracted with chloroform/methanol, deacylated in KOH/methanol, purified by preparative TLC and analyzed by Q-Exactive Orbitrap LC/MS as described in Methods.

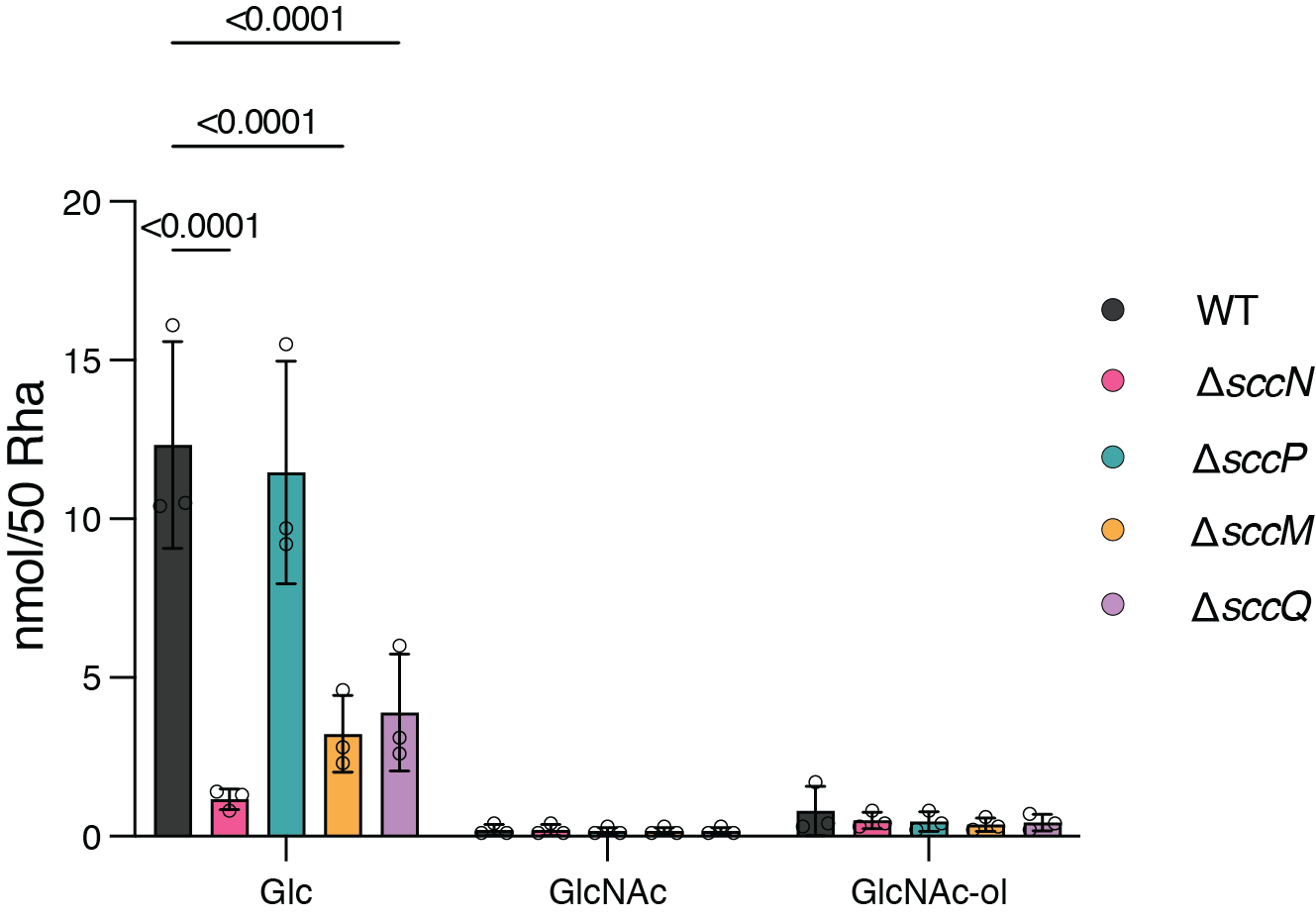

**Supplementary Fig. 4.** **Glycosyl content of SCCs derived from various deletion mutants of *S.*** ***mutans***

SCCs were released from cell wall preparations from the indicated mutant strains of *S. mutans* by mild acid hydrolysis, chemically reduced with sodium borohydride, partially purified by SEC on Biogel P150 and analyzed for glycosyl composition by gas chromatography/mass spectrometry as trimethylsilyl derivatives of *O*-methyl glycosides, as described in Methods. Moles of each sugar were normalized to a hypothetical polymer containing 50 Rha residues. Ordinary two-way ANOVA with Dunnett’s multiple comparison test was done for statistical analysis.

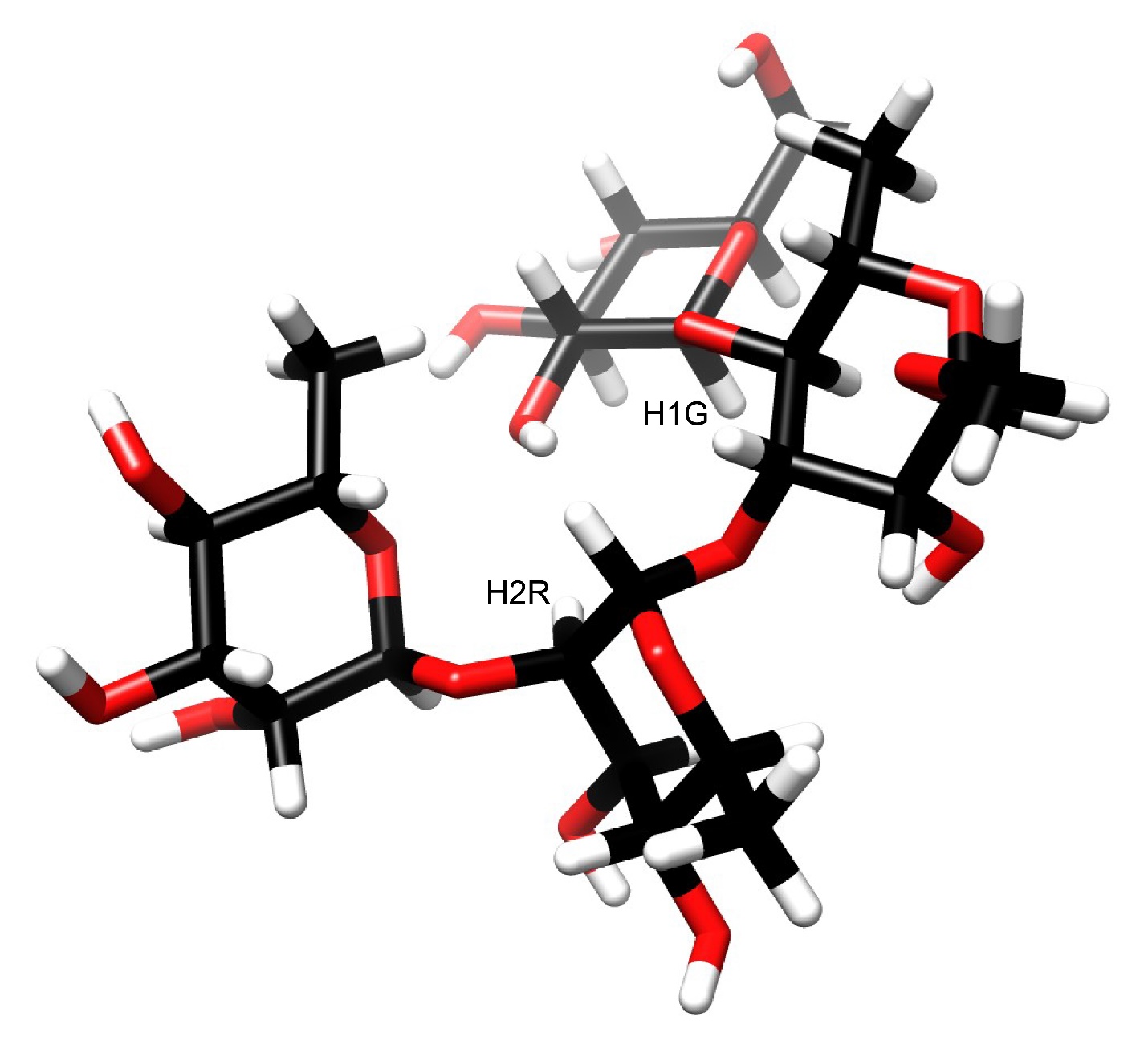

**Supplementary Fig. 5. Molecular model made by CarbBuilder ^10^ and graphically presented using UCSF Chimera ^11^ of the tetrasaccharide α-l-Rha*p*-(1→2)-α-l-Rha*p*-(1→3)[β-d-Glc*p*-(1→4)]-α-l-Rha*p*-OMe representing a branched region in the SCC.**

The anomeric proton (H1G) of the β-d-Glc*p*-(1→4)-linked side‑chain residue and the H2 proton (H2R) of the 2-linked rhamnosyl residue in the backbone show a mutual NOE in ^1^H,^1^H-NOESY NMR spectrum of the SCC due to spatial proximity, which is substantiated by the model.

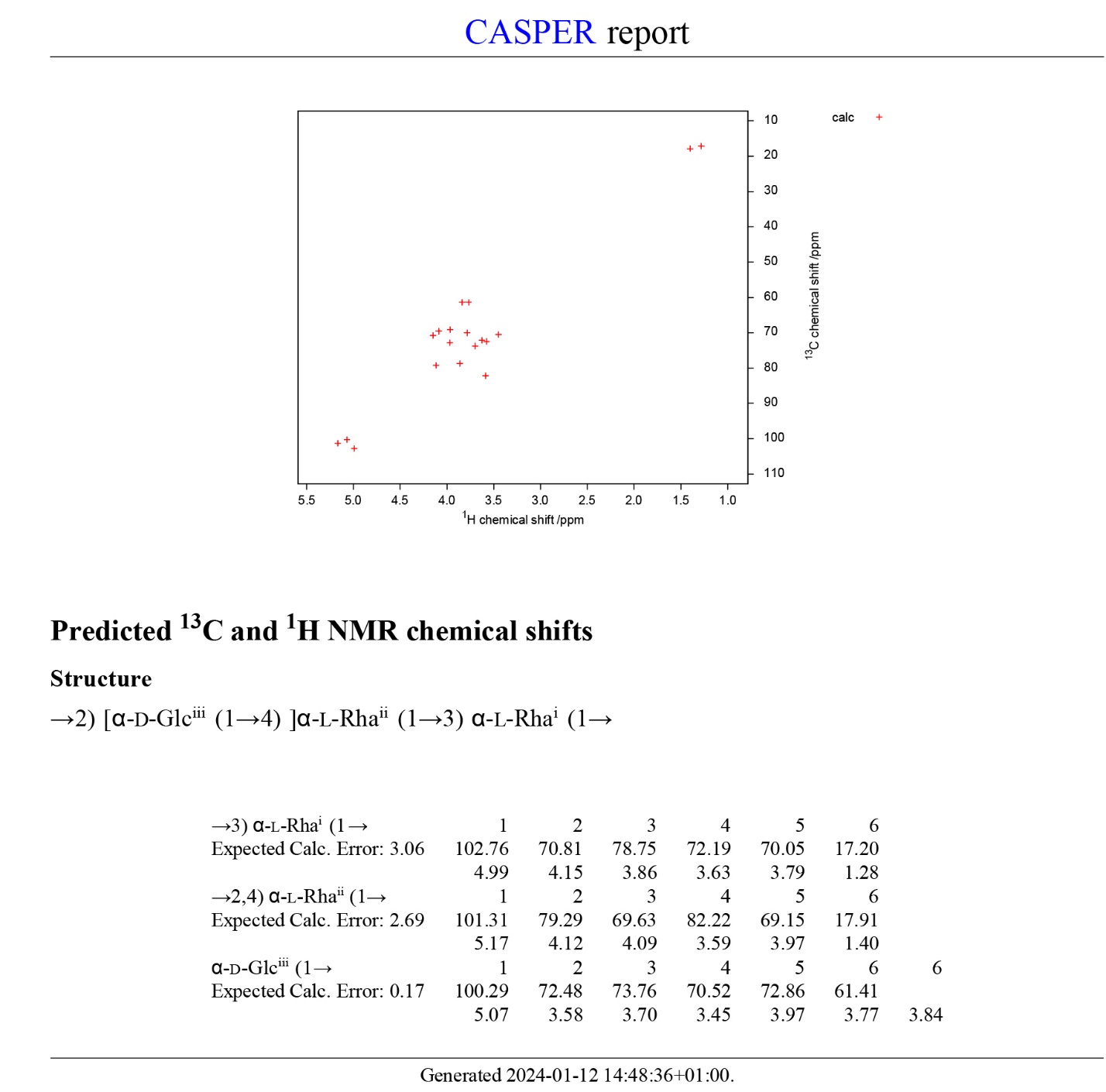

**Supplementary Fig. 6**. **NMR chemical shift prediction by CASPER ^12^ of the repeating unit structure containing the α-d-Glc*p*-(1→4)-linked side‑chain residue, detected in the ∆*sccM* SCC.**

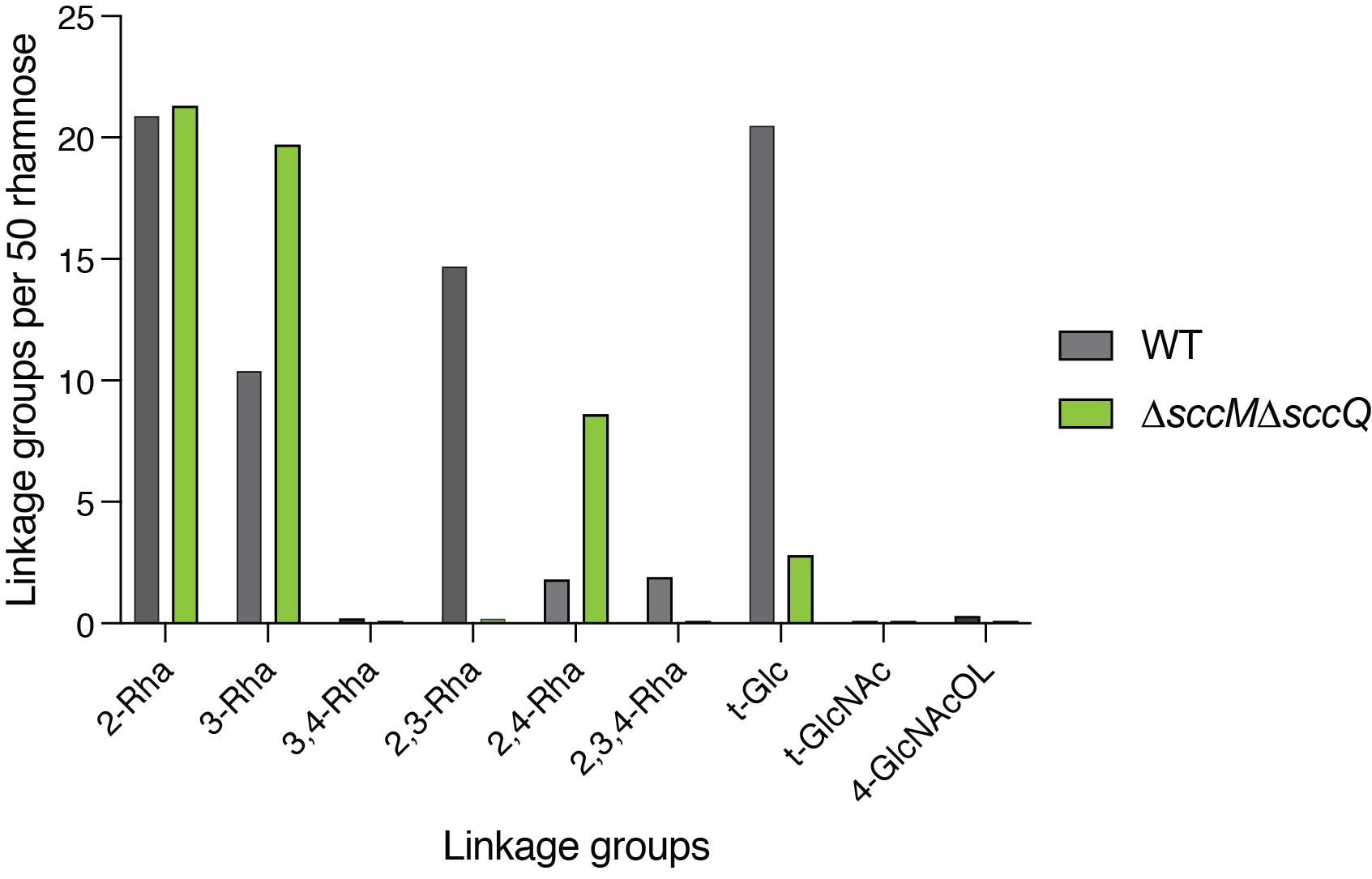

**Supplementary Fig. 7. Glycosyl linkage analysis of the WT and Δ*sccM*Δ*sccQ* SCCs**

Quantitative comparison of linkage groups detected in SCCs purified from WT and ∆*sccM*∆*sccQ* cell walls*.* Moles of each linkage type per molecule were calculated by multiplying the mol % for each component sugar by the relative area percent for each linkage type found in the gas chromatogram and then normalized to a hypothetical polymer containing 50 Rha units. Note that the only branched Rha unit detected in Δ*sccM*Δ*sccQ* SCC is 2,4-Rha.

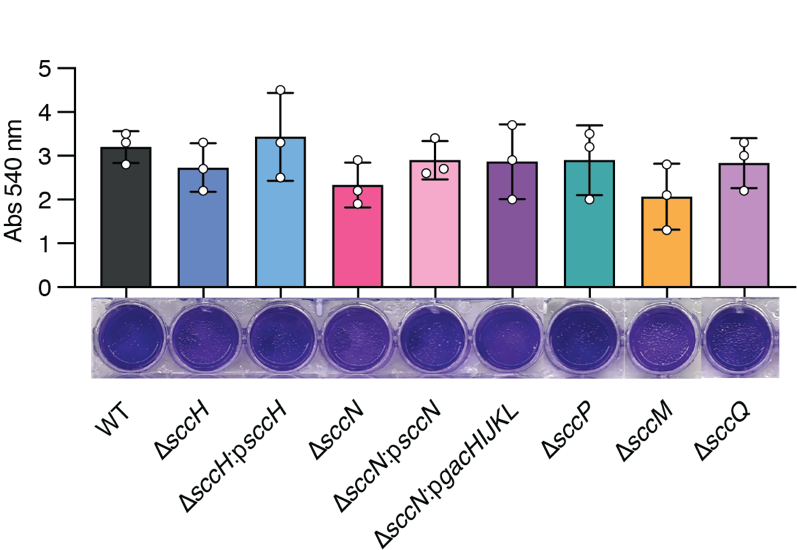

**Supplementary Fig. 8.** **Exopolysaccharide-based biofilms produced by *S. mutans* strains**

To obtain biofilms, *S.* *mutans* strains were incubated in UFTYE medium supplemented with 1% (wt/vol) sucrose for 24 h at 37 °C in presence of 5% CO_2_. Biofilms were analyzed as outlined in Methods. Image of crystal violet stained-biofilms (bottom panel) is a representative image of three independent experiments. Quantification of biofilm formation is shown in top panel. Columns and error bars represent the mean and S.D., respectively (n = 3). Data were analyzed using one-way ANOVA with Dunnett's multiple comparisons test. No significant differences were observed.

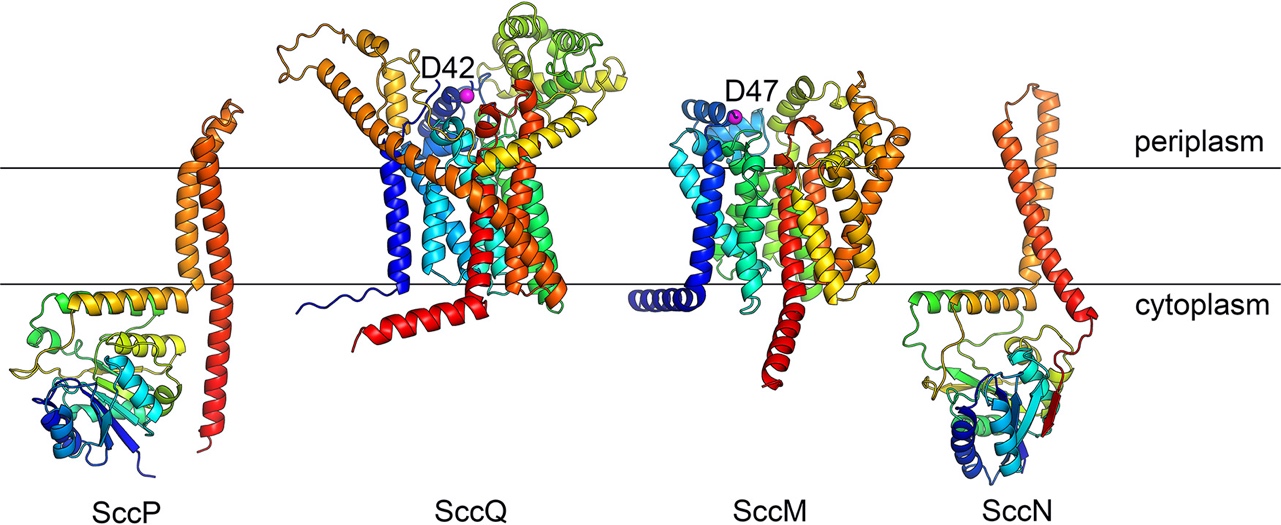

**Supplementary Fig. 9. Proposed topological orientation of glycosyltransferases involved in the glucosylation of *S.* *mutans* SCC**

Structural models predicted by AlphaFold2 pipeline ^13, 14^ for *S.* *mutans* SccP, SccQ, SccN and SccM are shown as cartoon diagrams colored in rainbow colors from N-terminus (blue) to C-terminus (red), relative to the bacterial plasma membrane. Magenta spheres highlight the key active-site residues of SccQ and SccM. The membrane topology was predicted using TopCons ^15^
